# Sensbio: An online server for biosensor design

**DOI:** 10.1101/2022.07.28.501596

**Authors:** Jonathan Tellechea-Luzardo, Raul Moreno López, Pablo Carbonell

**Affiliations:** Institute of Industrial Control Systems and Computing (AI2), Universitat Politècnica de València (UPV), 46022 Valencia, Spain

## Abstract

Allosteric transcription factor (aTF) based biosensors can be used to engineer genetic circuits for a wide range of applications. The literature and online databases contain hundreds of experimentally validated molecule-TF pairs, however the knowledge is scattered and often incomplete. Additionally, compared to the number of compounds that can potentially be produced using living systems, the number of those discovered with TF-based interactions is low. For these reasons, new tools that help researchers find new possible TF-ligand pairs are called for. In this work we present Sensbio (https://bit.ly/3OF4msH), an online web server that through similarity comparison against a TF-ligand reference database, is able to identify putative transcription factors that may be activated by a given input molecule. As a complement to the application, a predictive model has been developed to find new possible matches based on machine learning.

## Introduction

Biosensors allow researchers from various fields to use biological systems to detect external or internal signals and react in a designed manner [1]. Among other inputs, biosensors can be used to detect small molecules that may play important roles in bioremediation, metabolic engineering or biocomputing among other areas. An important class of biosensors are those based on allosteric transcription factors (aTFs) that bind to the signal molecule, triggering the expression or repression of a particular gene (e.g. a reporter gene). Even though biosensors have been used for a wide range of applications [2], the number of known responsive TFs is limited compared to the number of potential chemical targets.

During recent years both mining and experimental assays have been reported in the literature describing different methodologies to discover with new TF-ligand interactions. The whole procedure is a multi-step process that may, collectively, require years of research. The effort required to mine a new TF from the available genomic knowledge, to properly characterize it and to validate its functionality against a new molecule presents a high toll to pay for the biosensor designer. For these reasons, more computational databases and tools are needed to help in the design of new biosensors, specially in the prototype phase.

Here we present Sensbio, an easy-to-use web application that finds new possible TF-ligand interactions by protein sequence and molecular similarity analysis that can be additionally assisted by machine learning-based recommendations. The Sensbio open-source web server provides a set of tools to help in the design of transcription-factor based biosensor circuits. Based on a dataset containing 451 chemical compounds and 3507 transcription factor sequences, Sensbio assists synthetic biologists by suggesting potential new TF-ligand interactions based on 6 different sources of transcription factor data, finding similar molecules and candidate transcription factors to the inputs. The Sensbio web server allows users to identify existing and novel transcription factor-based biosensors for applications ranging from genetic circuits design, screening, production and bioremediation of chemicals to diagnostics.

## Material and methods

### Databases, packages and tools used in this study

The dataset published by Koch et al. [3] was used as a starting point for the Sensbio database. It contains a 2018 collection of TF-ligand interactions from different databases and literary resources. To expand and update this dataset, data dumps detailing aTFs and their triggering compounds were collected, cleaned and formatted accordingly from the following databases: BioNemo [4], RegulonDB [5], RegPrecise [6], RegTransBase [7], Sigmol [8] and GroovDB (previously gBiosensor) [9].

Custom Python 3 scripts (using standard libraries like Pandas and Numpy) were used to populate, clean, format and analyze the database and to build the web application through the Streamlit framework (https://streamlit.io/). Molecular fingerprints were extracted, analyzed and compared using the RDKit python library [10]. Networkx python module was used to describe and produce the molecular network. A local BLAST+ installation allowed the scoring and ranking of the protein sequences. Ete3 python toolkit produced the phylogenetic trees of the TF sequences. Deep learning techniques were applied to build the predictive model through the Tensorflow and Keras python libraries.

Classyfire [11] and iFragment [12] external web applications were used to classify the different molecules by chemical and metabolic categories respectively. Classyfire produces a hirearchical list of ontologies. In this case, the parent ontology was kept as the final category for each molecule. iFragment on the other hand, produces a list of KEGG [13] metabolic pathways ordered by the probability of the input compound to belong to that particular pathway. The three pathways with the lowest p-value were selected. Using the KEGG restful API (https://www.kegg.jp/kegg/rest/keggapi.html), the parent ontology was extracted for each pathway and assigned as the final metabolic category.

### Implementation

First, the Sensbio database was built detailing both molecular (molecule common name, SMILES, InChi and information on the metabolic paths where the molecule plays a role) and protein sequence information (TF name, origin species, protein sequence, NCBI and Uniprot accession numbers and database and literature references) for each of the TF-ligand pairs mined from the previously detailed databases and bibliographic sources.

Sensbio accepts protein sequences and chemical compounds as inputs. Two possible use cases for the application are envisioned:

#### Use case 1

When the user wants to determine if a chemical compound can be sensed using TF-based biosensors they can use the molecular similarity tool (Figure 1, red flow). This approach is based on the RDkit library to calculate a Tanimoto similarity score [14] of the input molecule against all the other compounds in the database. As output, Sensbio provides a set of ranked entries similar to the input chemical structure.

**Figure 1.**
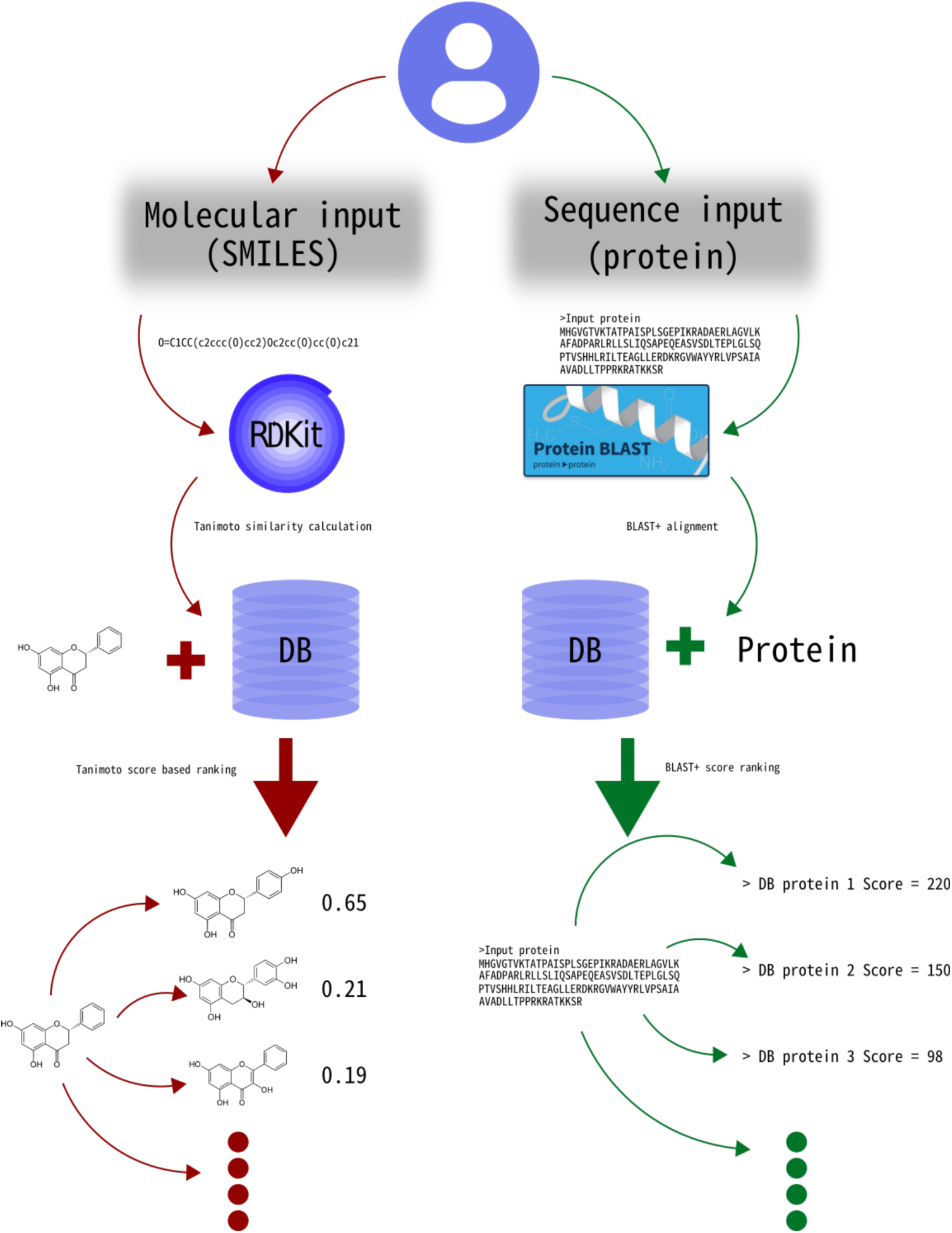
Sensbio workflows. Red flow: a molecular input by the user produces an ordered rank of similar molecules paired with the aTF that is activated or repressed by them. Green flow: a protein sequence input produces a ranked list of sequences and their binding molecule.

#### Use case 2

When the user wants to check a predicted putative TF for sensing capabilities they can use the sequence similarity tool (Figure 1, green flow). This tool is based on BLAST+ algorithm and, using the user’s input as a query, it provides the top significant set of entries in the database closest to the input sequence as output. This can be used to determine possible molecular ligands and to fast-track a literature search on the closest transcription factors.

Apart from these use cases, the application can be used to visualize the state-of-the-art knowledge of the aTF-mediated biosensing space. The repository containing all the application files and requirements is available at: https://github.com/jonathan-tellechea/sensbio_app.

##### Predictive model

Moreover, a predictive system has been developed with the aim of having a machine-learning base recommendation system for finding new possible TF-ligand interactions. In order to train the model, the Sensbio database was initially used. For the TF sequences, the one-hot encoding technique was used. For the molecules, fingerprints from SMILES were extracted. Also, negative cases, i.e., cases where there is no affinity between the TF and the molecule were generated. For this purpose, a molecule that does not resemble the molecule associated with a given TF based on their Tanimoto index was randomly selected for each sequence.

The network architecture (Figure 2) is based on two branches (one for each type of input), which are then concatenated. For the TF branch a LSTM layer was considered as it can learn from sequential data [15].

**Figure 2.**
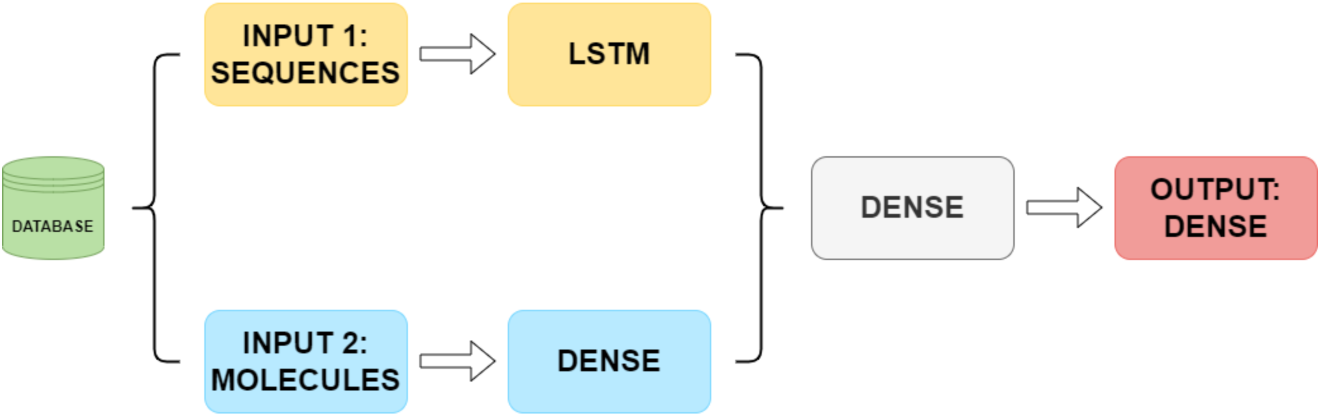
Network architecture diagram

In terms of model training, the K-Fold Cross-Validation technique was carried out to test the different possibilities of the hyperparameters of the model. The model returns a score between 0 and 1, where the lowest value indicates that there is not affinity between the TF and the molecule, and the highest indicates the opposite. The repository containing the codes required to build and train this model are available at: https://github.com/mlraul/biosensor_predictor.

## Results

### Molecular similarity

Next it is described the expected results of the molecular similarity tool. For this purpose, naringenin (O=C1CC(c2ccc(O)cc2)Oc2cc(O)cc(O)c21) and pinocembrin (C1C(OC2=CC(=CC(=C2C1=O)O)O)C3=CC=CC=C3) molecules are used as examples. When naringenin is fed in the chemical tool, it produces the dataset shown in Table 1. The Tanimoto score of 1 for the first 5 entries confirms that the database contains the exact same molecule and provides information on 5 TFs that have been described to be activated by the compound. This informs the biosensor designer that the input molecule has been described as the activator of these TFs so they can make a decision on their following experimental workflow.

**Table 1.**
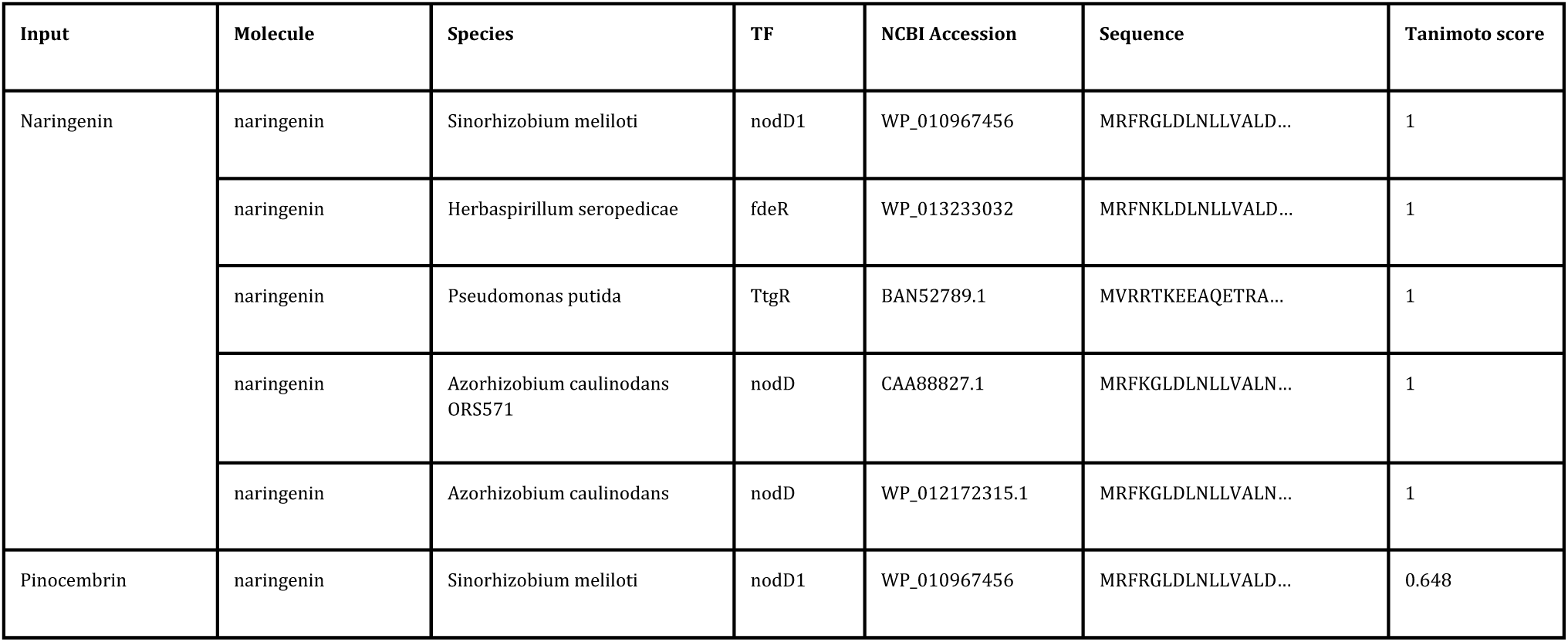
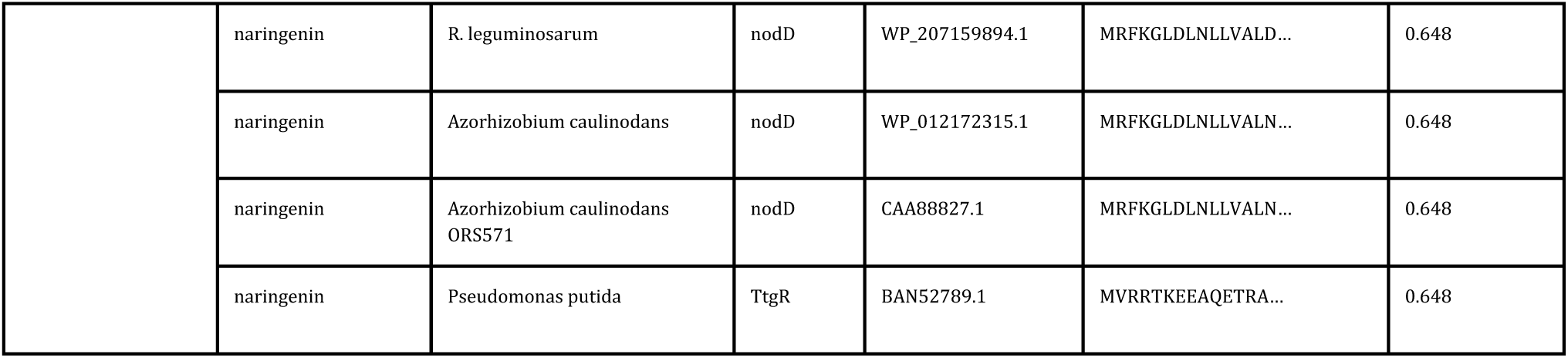
Sensbio example molecular results.

When a molecule that is not in the database is provided as input, the tool provides the set of entries ordered by Tanimoto score. In the case of the pinocembrin, the application ranks highest naringenin entries by close similarity to the compound, suggesting that pinocembrin could be sensed though naringenin-activated TFs. This was experimentally confirmed in Trabelsi et al [16]. This information could be used by the user to find TF that are likely to sense their input compound and build prototype biosensor circuits around this information.

### Sequence similarity

Here we showcase the behavior of the application when using its sequence similarity feature. Given a TF sequence that is present in the database (e.g. AseR, *B. subtilis*, NP_388414.1 which is triggered by arsenite) the application produces the ranked entries shown in Table 2 (a summarized view of the whole output data). In essence, the software recognizes the sequence as present in the database by giving it the highest rank (based on the BLAST+ scoring system) and 100% identity score. It also provides the user with other relevant sequences that recognize the same compound that may be worth studying further for increased biosensor design space in the laboratory.

**Table 2.**
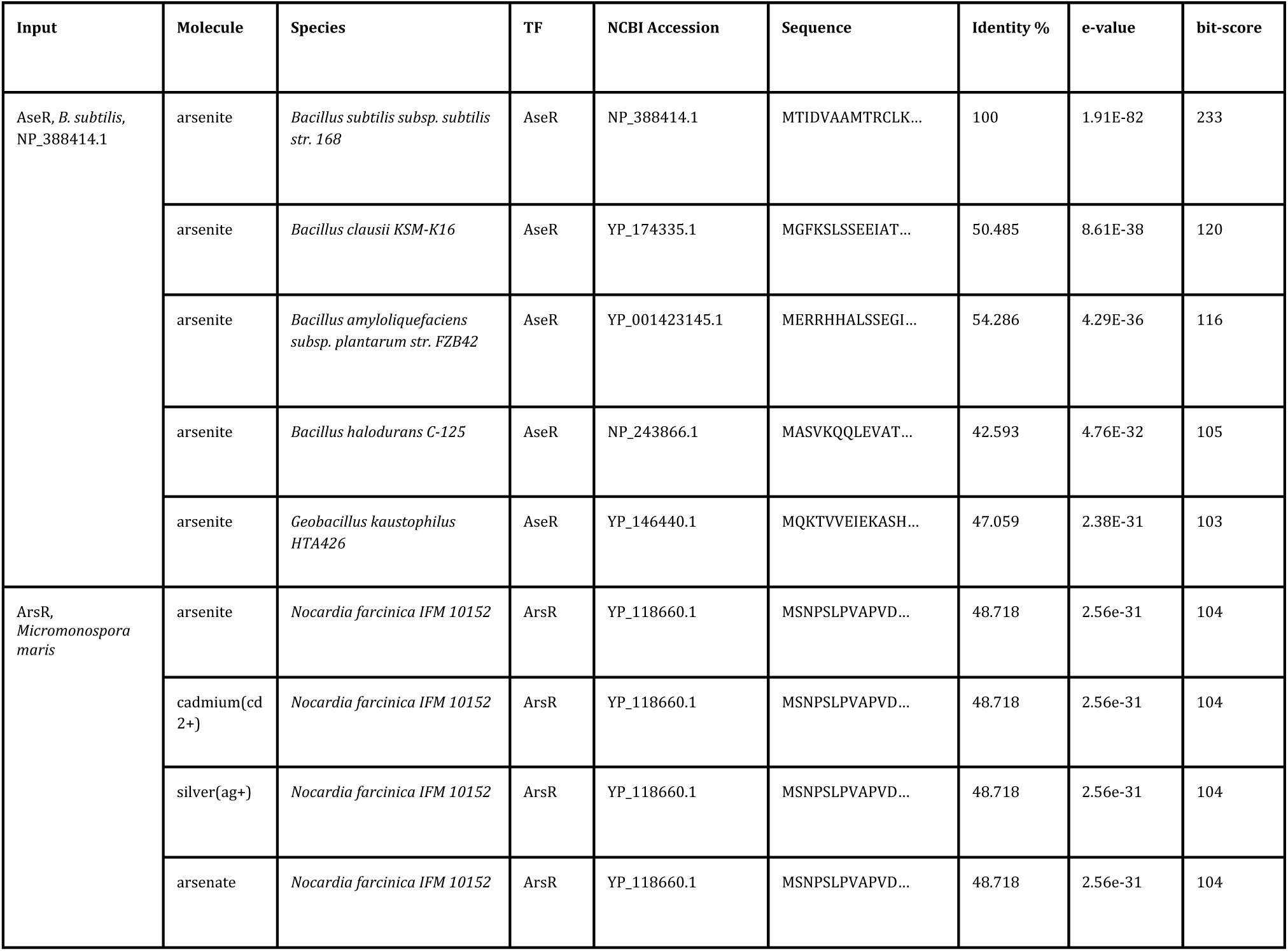
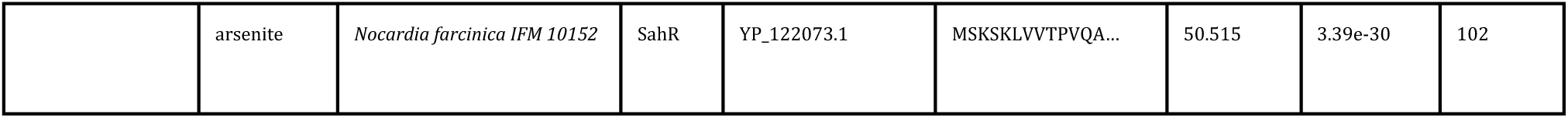
Sensbio example sequence results.

When a different TF, not present in the database, is fed to the sequence similarity tool (e.g. ArsR, *Micromonospora maris*, WP_043720559.1) one should expect the results shown in Table 2. Again, the script returns a list of the most similar proteins in the database together with information on the species and triggering molecules. This information could be used after discovering a new TF to assess possible molecular targets together with other sources of information before experimental validation.

### Database analysis

Finally, we highlight in this section the most important features of the Sensbio database collected as previously described. It contains 451 unique molecules and 3507 protein sequences which interact among themselves producing 5387 unique TF-ligand pairs.

Using the RDkit python library, the Tanimoto score of all the molecules against each other was calculated. This similarity matrix can be used to assess how similar are the molecules of the database (Figure 3). These results show that most of the molecules are very similar (Tanimoto scores between 0.9 and 1.0).

**Figure 3.**
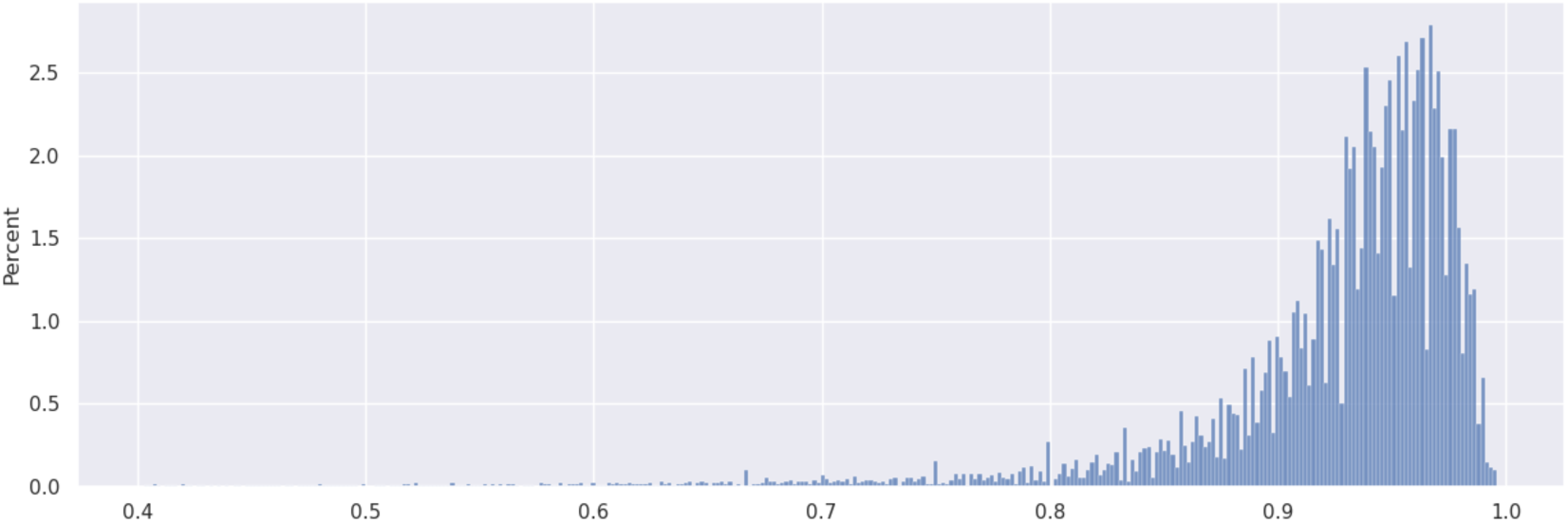
Tanimoto score distribution of the whole molecular collection (451 molecules).

Further analysis using the network python library NetworkX shows how the molecules are related and clustered together by the similarity score (Figure 4). The network figure confirms the similarity distribution clustering the molecules in 4-5 well-defined molecule categories.

**Figure 4.**
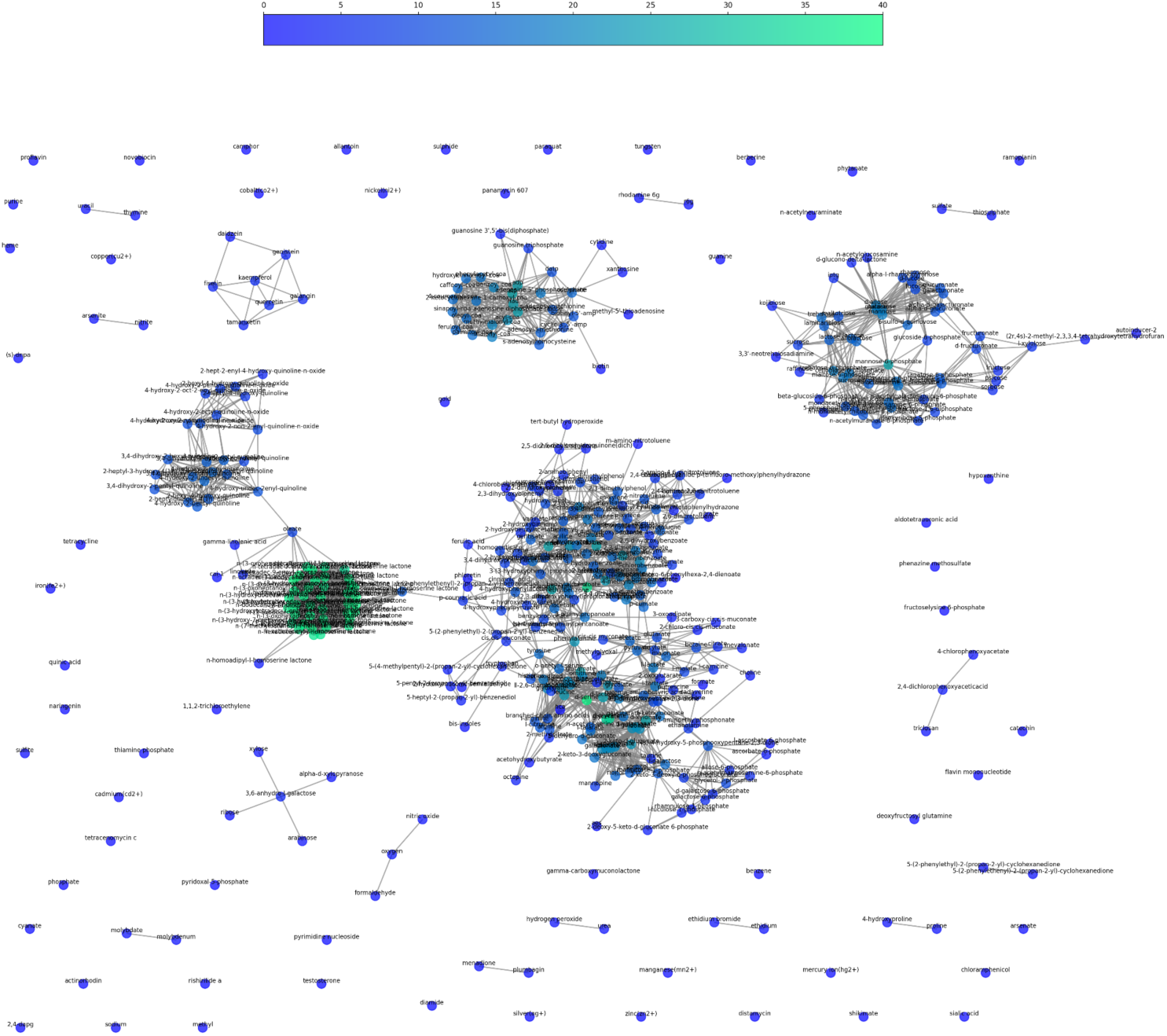
Molecular network and clustering of the database. Two molecules are connected together if they have a Tanimoto higher than 0.75. The color of the node represent the number of connections of that node.

The molecules can be classified using different criteria. First, molecules were classified using chemical ontologies using the Classyfire tool. Figure 5 shows the different chemical categories present in the database and their abundance. The most common category was established as “Hydrocarbon derivatives” (simple and complex sugars, etc.), followed by “Carbonyl compounds” (some amino acids, lactones, etc.).

**Figure 5.**
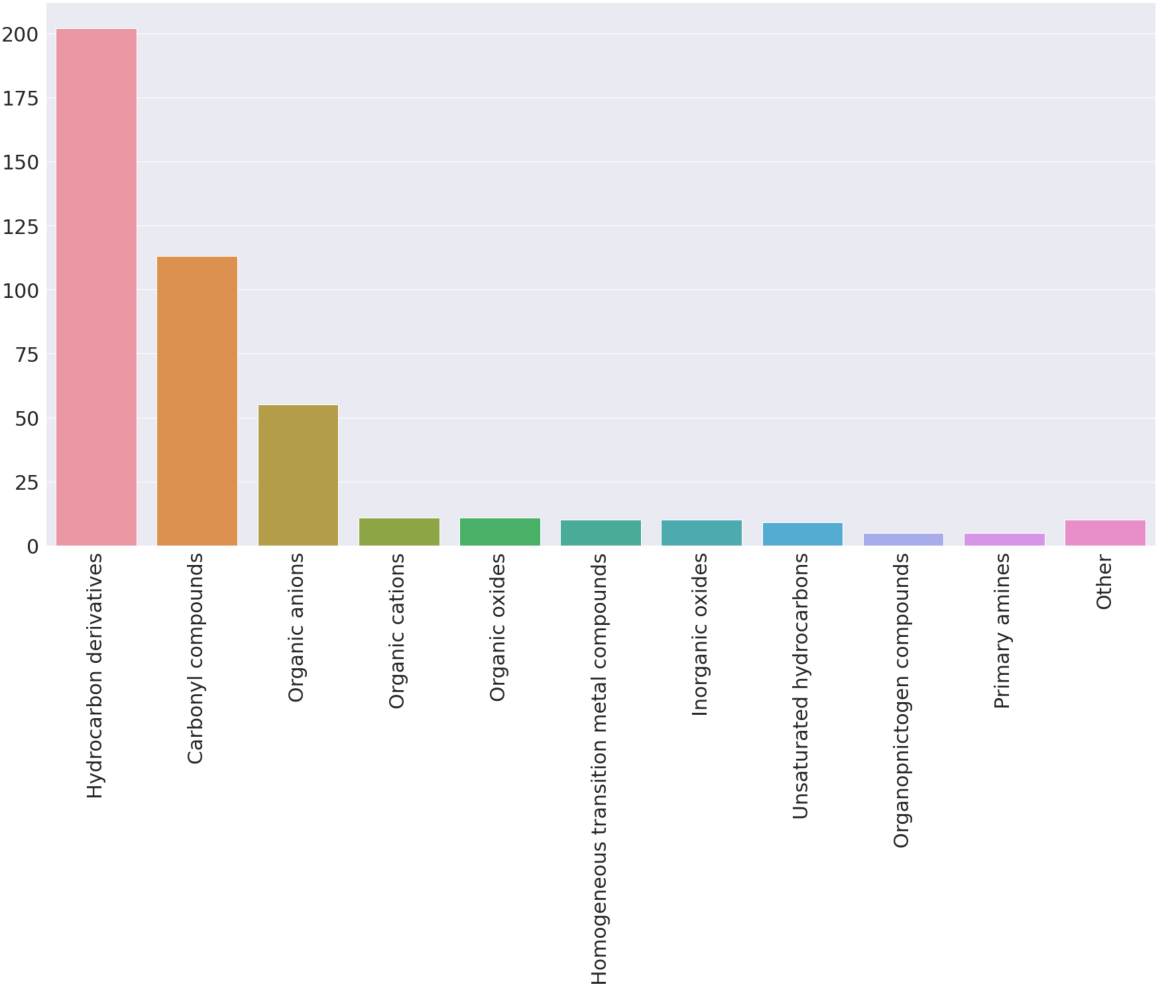
Frequency of molecular ontologies discovered in the database. A total of 25 molecular categories are present.

Another classification can be made using metabolism as main criteria. The iFragment tool was used to assign biological pathways to each of the compounds in the database searching against the KEGG pathways dataset. Figure 6 shows the distribution of different KEGG pathways found. Note that the three most likely KEGG functions assigned to each compound were kept. Most of the molecules in the dataset are related to amino acid metabolism.

**Figure 6.**
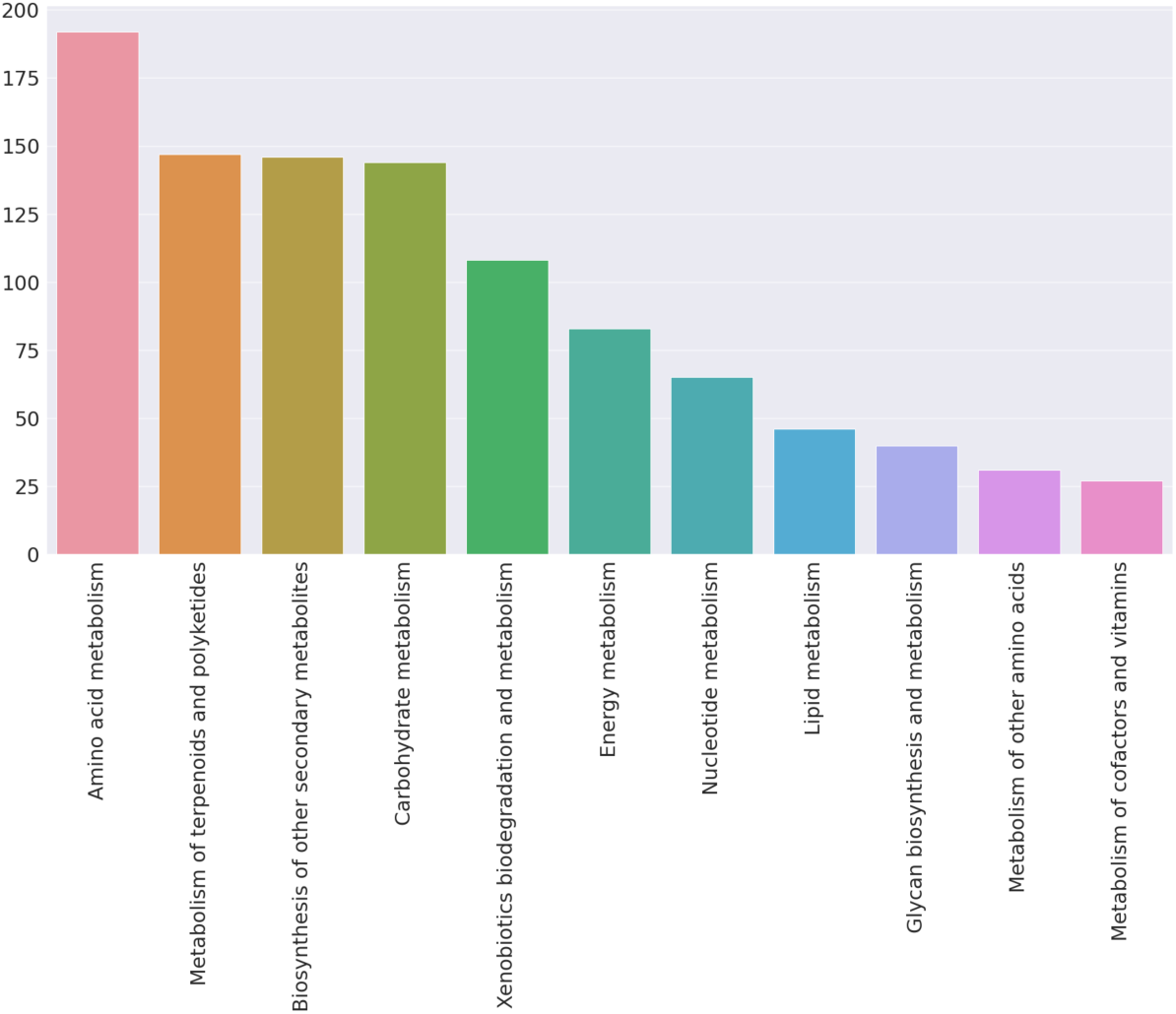
Frequency of metabolic categories found in the database.

The protein sequences in the database were analyzed by their relationship with their compound pair. The table in Figure 7 details how the sequences are related to the chemical categories.

**Figure 7.**
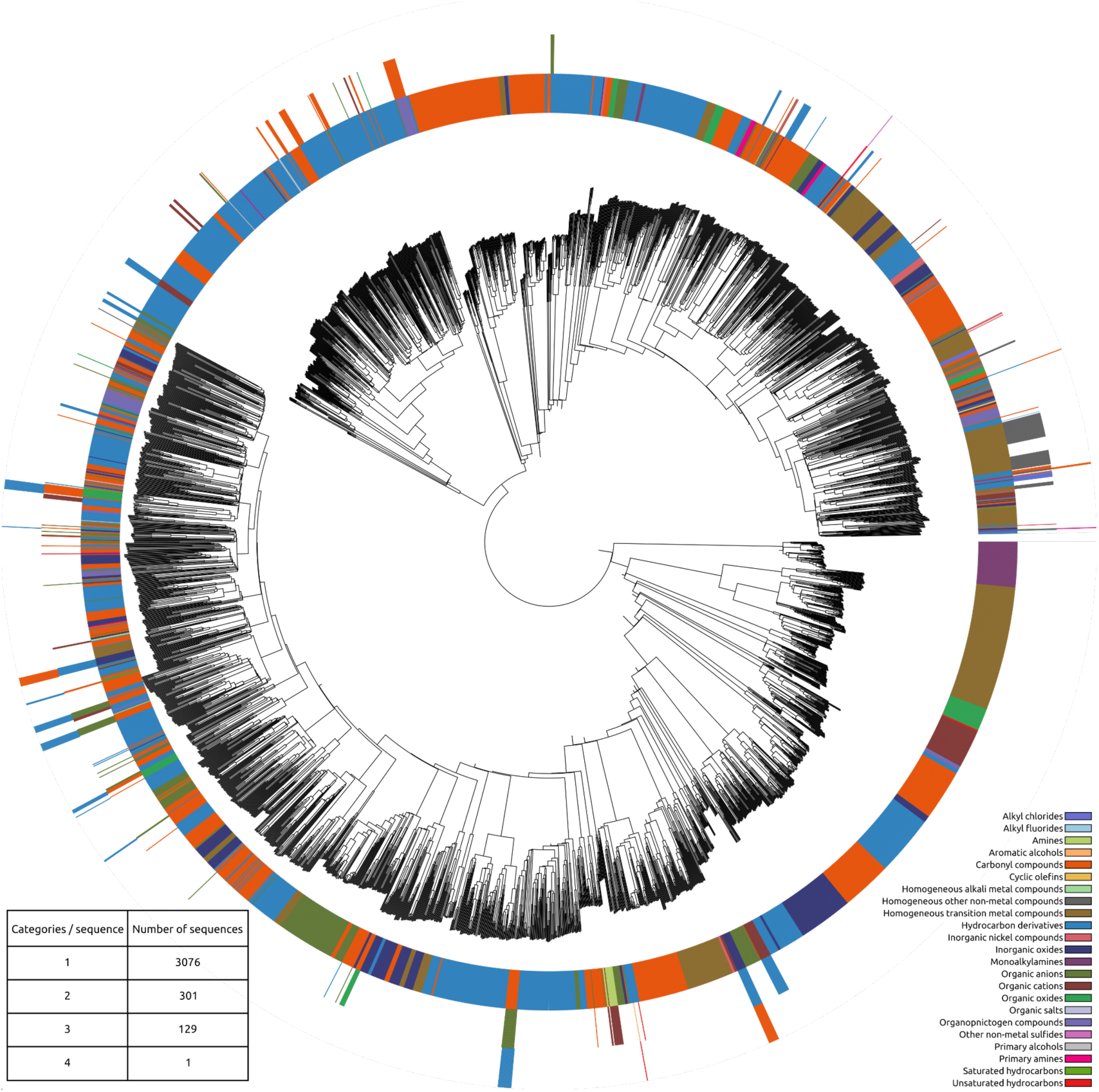
Phylogenetic tree of the protein sequences in the database paired with the molecular categories of the compound(s) that bind to the TF.

87.7% of the 3507 unique sequences have been associated to a single molecular ontology. The rest are “promiscuous” TFs and are triggered by more than one molecular category.

The 3507 sequences were aligned and assembled in a phylogenetic tree using Clustal Omega. The ete3 python library was used to produce the tree figures coupled with the categorical information. The chemical and metabolic categories previously determined were paired with each sequence in the tree producing the Figure 7 and Figure 8.

**Figure 8.**
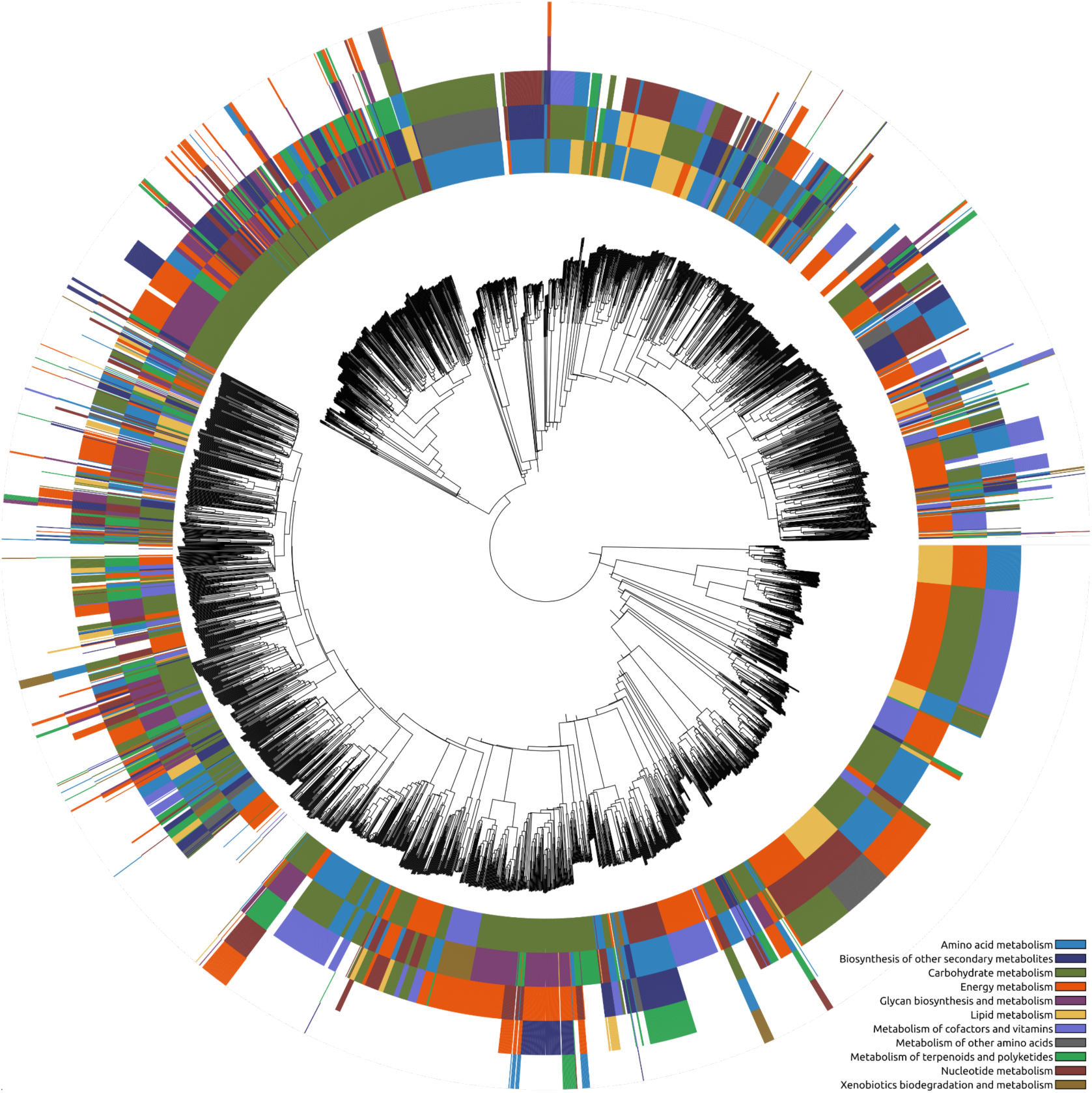
Phylogenetic tree of the protein sequences in the database paired with the metabolic categories of the compound(s) that bind to the TF.

### Predictive model performance

For the machine learning-based model shown in Figure 2, loss and accuracy metrics were used during the validation process. Their evolution curves over the epochs of training during the last validation are shown in Figures 9 and 10. The average loss value was 0.4 and the accuracy value was 88.2%.

**Figure 9.**
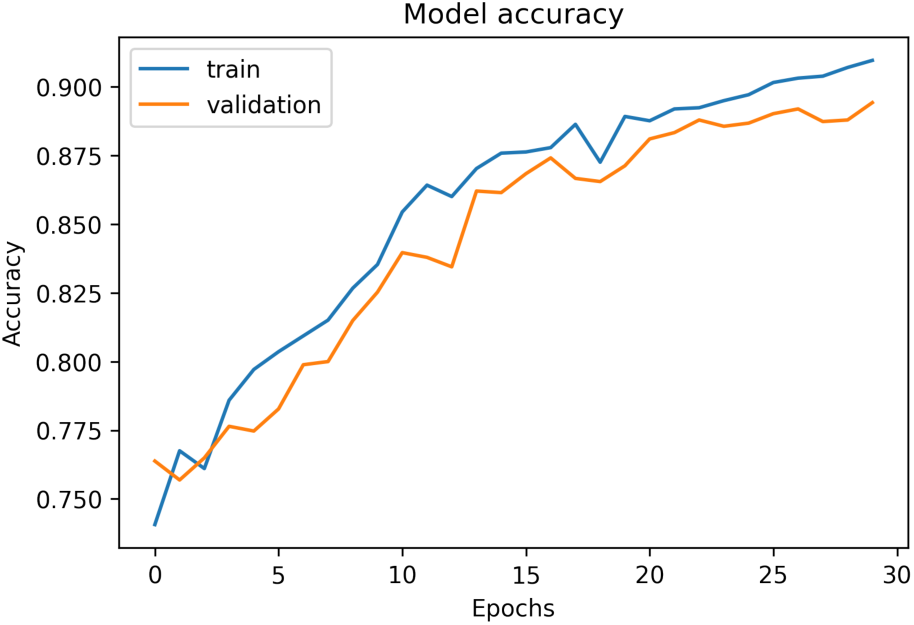
Example of accuracy curve for one of the validation processes of the final model.

**Figure 10.**
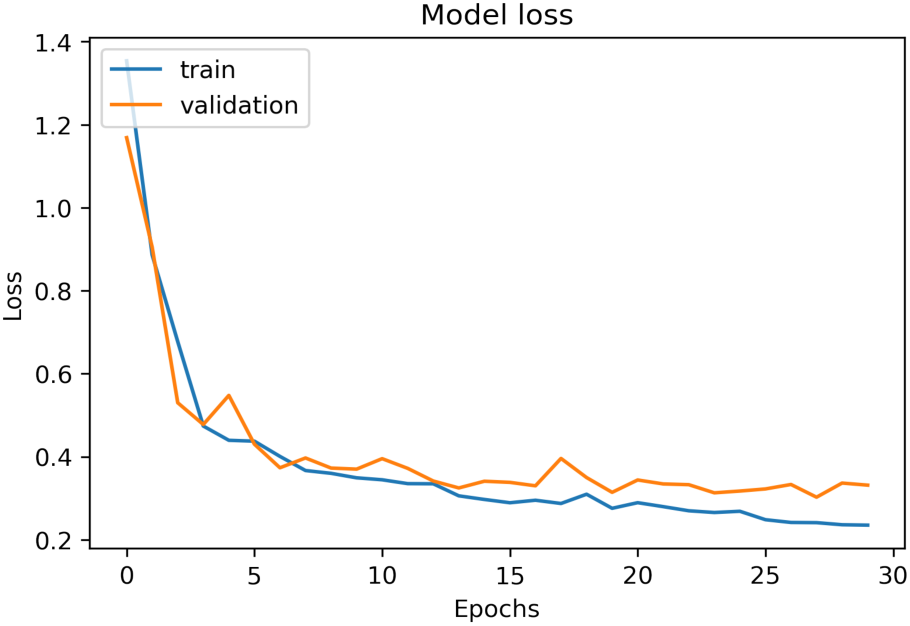
Example of loss curve for one of the validation processes of the final model.

The stabilization of both, accuracy and loss curves, and the evolution of the validation curve with respect to the train one, show that the number of epochs is enough to obtain acceptable results and that there is no overfitting.

After the validation, the actual model training was carried out. The scores obtained when predictions were made with the test data have been 0.3 of loss and 89.8% of accuracy. The ROC curve (Figure 11) and the AUC value have also been obtained.

**Figure 11.**
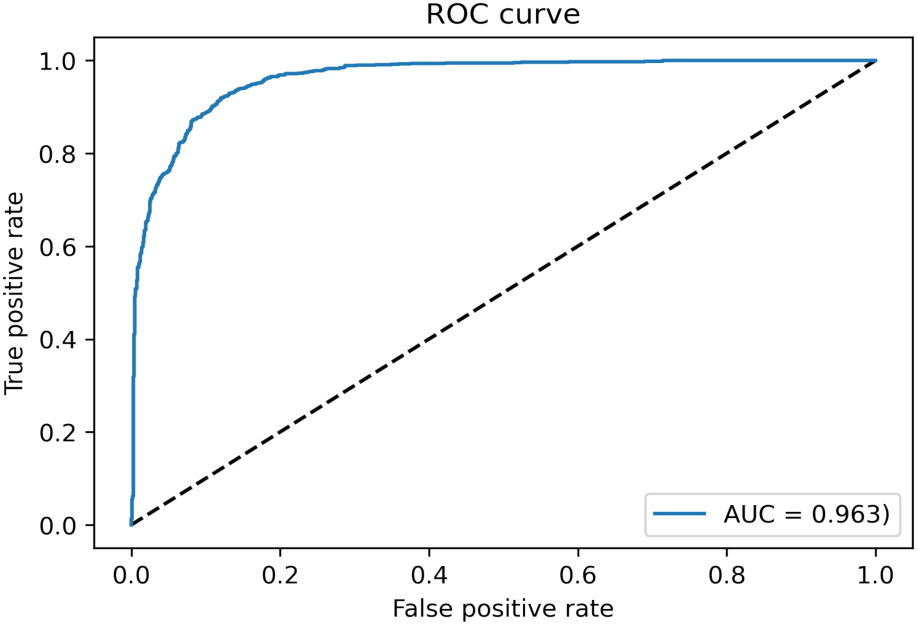
ROC curve resulting from test and its AUC value

The evolution of the ROC curve and the AUC value associated led to the conclusion that the model performs reasonably well when distinguishing between positive (there is affinity between the TF and the ligand) and negative (there is no affinity) cases.

Lastly, when comparing performance for each class (affinity between the TF and the molecule or not), a F1-score close to 0.9 has been obtained for both positive and negative cases. The similarity between the F1-score of the two groups demonstrates that the model is well balanced for predicting either the affinity between a TF and a molecule or the impossibility of using the TF to sense the molecule.

### Conclusions and future directions

In this study we present two resources that may ease the biosensor design process and help researchers prototype biosensing circuits faster.

The first one is the web server application called Sensbio. This easy-to-use application can suggest putative aTFs that may be able to detect a given input compound. The tool can also be used to determine the possible ligand molecule of a newly discovered TF sequence by homology to the database. The tool is available at https://bit.ly/3OF4msH.

Secondly, the ML model produced in this study can be used to find extra TF-ligand interactions based on AI. Even if results are promising, predictions of the ML-based model still lack enough specificity, as we are expecting to use this tool in order to refine the homology search. Future work will test other model architectures, including using the homology search results as additional input to the model.

Besides the improvement of ML-based predictions, the current dataset can be augmented with protein homologues to improve further the prediction metrics. In the future, the ML model will be improved and integrated in the application. This could add an extra layer of certainty to trust the predicted TF-ligand interaction based on other factors than sequence or molecular similarity. An additional layer of information useful to the users may be the computation of structural-based scores for each TF-ligand pair from tools like molecular docking. Finally, at some point we would like to move the application to a more customizable and stable location (either local or in-the-cloud).

